# Uneven distribution of cobamide biosynthesis and dependence in bacteria predicted by comparative genomics

**DOI:** 10.1101/342006

**Authors:** Amanda N. Shelton, Erica C. Seth, Kenny C. Mok, Andrew W. Han, Samantha N. Jackson, David R. Haft, Michiko E. Taga

## Abstract

The vitamin B_12_ family of cofactors known as cobamides are essential for a variety of microbial metabolisms. We used comparative genomics of 11,000 bacterial species to analyze the extent and distribution of cobamide production and use across bacteria. We find that 86% of bacteria in this data set have at least one of 15 cobamide-dependent enzyme families, yet only 37% are predicted to synthesize cobamides *de novo*. The distribution of cobamide biosynthesis varies at the phylum level, with 57% of Actinobacteria, 45% of Proteobacteria, and 30% of Firmicutes, and less than 1% of Bacteroidetes containing the complete biosynthetic pathway. Cobamide structure could be predicted for 58% of cobamide-producing species, based on the presence of signature lower ligand biosynthesis and attachment genes. Our predictions also revealed that 17% of bacteria that have partial biosynthetic pathways, yet have the potential to salvage cobamide precursors. These include a newly defined, experimentally verified category of bacteria lacking the first step in the biosynthesis pathway. These predictions highlight the importance of cobamide and cobamide precursor crossfeeding as examples of nutritional dependencies in bacteria.

## Introduction

Microorganisms almost universally reside in complex communities where individual members interact with each other through physical and chemical networks. A major type of chemical interaction is nutrient crossfeeding, in which microbes that lack the ability to synthesize particular required nutrients (termed auxotrophs) obtain these nutrients from other organisms in their community (Seth and Taga, 2014). By understanding which organisms require nutrients and which can produce them, we can predict specific metabolic interactions between members of a microbial community (Abreu and Taga, 2016). With the development of next-generation sequencing, the genome sequences of tens of thousands of bacteria from diverse environments are now available, leading to the possibility of predicting community interactions based on the genomes of individual members. However, the power to predict the metabolism of an organism by analyzing its genome remains limited.

The critical roles of cobamides (the vitamin B_12_ family of enzyme cofactors) in the metabolism of humans and diverse microbes has long been appreciated. Only recently, however, has cobamide-dependent metabolism been recognized as a potential mediator of microbial interactions (Degnan *et al.*, 2014b; Seth and Taga, 2014). Cobamides are used in a variety of enzymes in prokaryotes, including those involved in central metabolic processes such as carbon metabolism and the biosynthesis of methionine and deoxynucleotides (Fig. 1). Some of the functions carried out by cobamide-dependent pathways, such as acetogenesis via the Wood-Ljungdahl pathway in anaerobic environments, can be vital in shaping microbial communities (Ragsdale and Pierce, 2008). Cobamides are also used for processes that are important for human use, such as reductive dehalogenation and natural product synthesis (Banerjee and Ragsdale, 2003; Broderick *et al.*, 2014).

**Figure 1:**
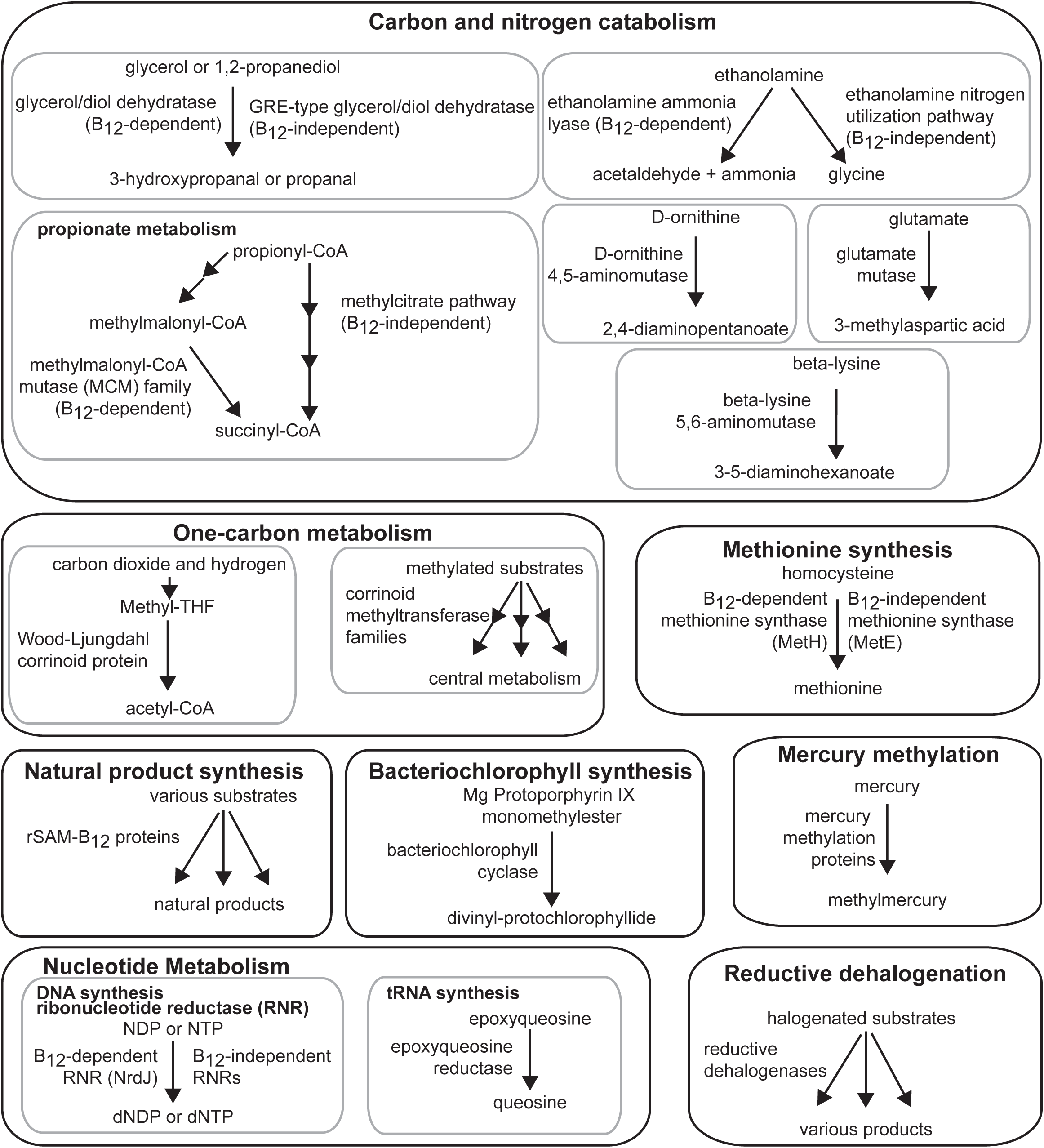
Functions carried out by cobamide-dependent processes. Reactions carried out by cobamide-dependent enzymes are shown on the left side of the arrows and cobamide-independent alternative processes, if known, on the right. Annotations or query genes used for searching for each function are listed in Supplementary Table 4.

*De novo* cobamide biosynthesis involves approximately 30 steps (Warren *et al.*, 2002), and the pathway can be divided into several segments (Fig. 2). The first segment, tetrapyrrole precursor biosynthesis, contains the first five steps of the pathway, most of which are also common to the biosynthesis of heme, chlorophyll, and other tetrapyrroles. The next segment, corrin ring biosynthesis, is divided into oxygen-sensitive (anaerobic) and oxygen-dependent (aerobic) routes, depending on the organism. These two alternative pathways then converge at a late intermediate, which is further modified to form the cobamide (Fig. 2, nucleotide loop assembly). The latter portion of the pathway involves adenosylation of the central cobalt ion followed by the synthesis and attachment of the aminopropanol linker and lower axial ligand (Fig. 2). Investigation of cobamide crossfeeding must account for structural diversity in the lower ligand (Fig. 2B), as only a subset of cobamide cofactors can support growth of any individual organism (Yan *et al.*, 2012; Mok and Taga, 2013; Degnan *et al.*, 2014a; Helliwell *et al.*, 2016; Keller *et al.*, 2018). Recent work has identified many of the genetic determinants for the biosynthesis of the benzimidazole class of lower ligands (Campbell *et al.*, 2006; Taga *et al.*, 2007; Gray and Escalante-Semerena, 2007; Hazra *et al.*, 2015; Mehta *et al.*, 2015) and attachment of phenolic lower ligands (Chan and Escalante-Semerena, 2011; Newmister *et al.*, 2012) (Fig. 2).

**Figure 2:**
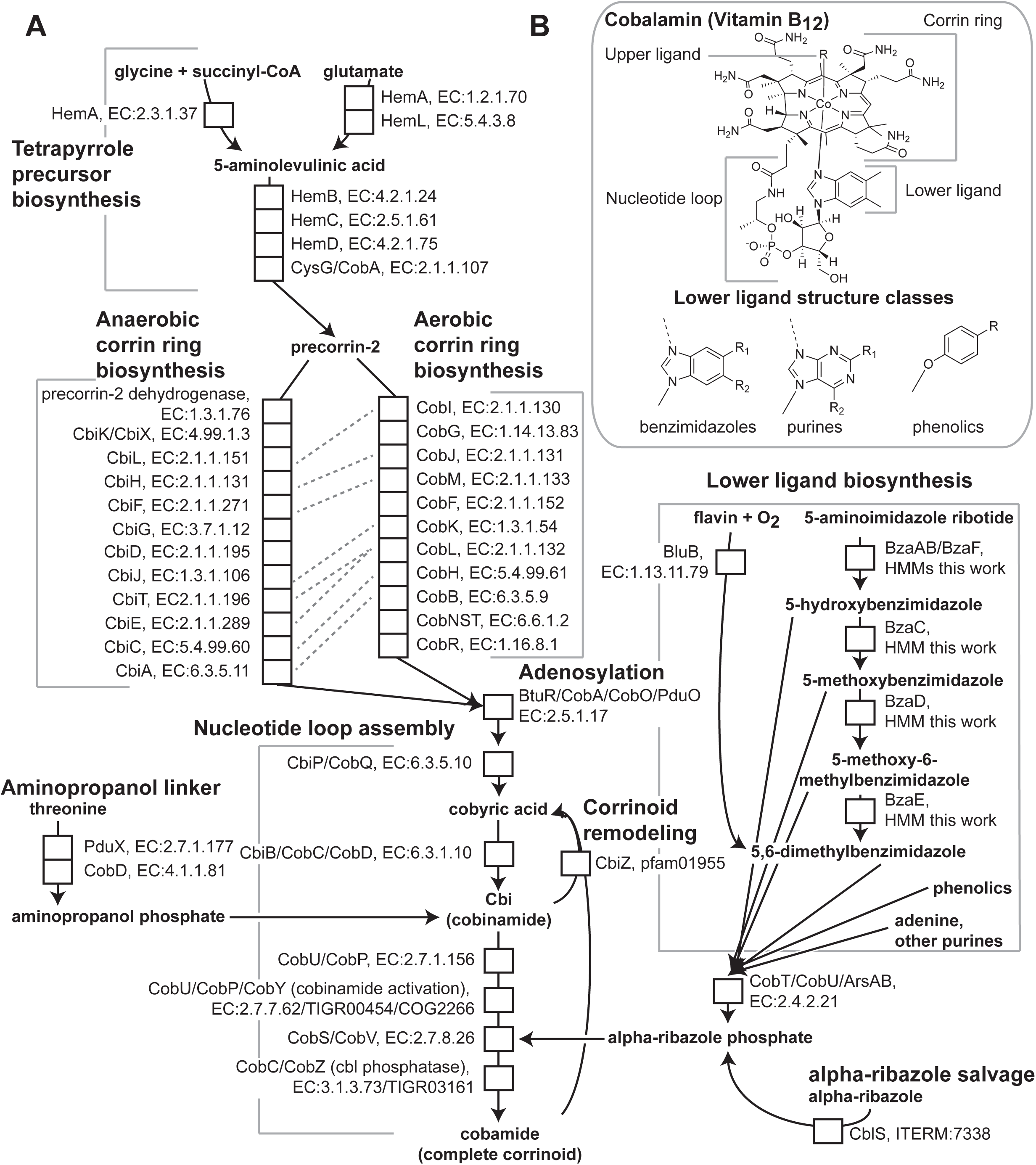
Cobamide biosynthesis and structure. **A.** The cobamide biosynthesis pathway is shown with each enzymatic step indicated by a white box labeled with the gene names and functional annotation. Subsections of the pathway and salvaging and remodeling pathways are bracketed or boxed with labels in bold. Orthologous enzymes that carry out similar reactions in aerobic and anaerobic corrin ring biosynthesis are indicated by dashed lines. **B.** Structure of cobalamin. The upper ligand R can be an adenosyl or methyl group. Classes of possible lower ligand structures are also shown. Benzimidazoles: R1=H, OH, CH_3_, OCH_3_; R2=H, OH, CH_3_, OCH_3_. Purines: R1=H, CH_3_, NH_2_; R2=H, NH_2_, OH, O. Phenolics: R=H, CH_3_.

Previous analyses of bacterial genomes have found that less than half to three fourths of prokaryotes that require cobamides are predicted to make them (Rodionov *et al.*, 2003; Zhang *et al.*, 2009), suggesting that cobamide crossfeeding may be widespread in microbial communities. Analyses of cobamide biosynthesis in the human gut (Degnan *et al.*, 2014a; Magnúsdóttir *et al.*, 2015) and in the phylum Cyanobacteria (Helliwell *et al.*, 2016) further reinforce that cobamide-producing and cobamide-dependent bacteria coexist in nature. These studies provide valuable insights into the extent of cobamide use and biosynthesis in bacteria, but are limited in the diversity and number of organisms studied and have limited prediction of cobamide structure.

Here, we have analyzed the genomes of over 11,000 bacterial species and generated predictions of cobamide biosynthesis, dependence, and structure. We predict that 86% of sequenced bacteria are capable of using cobamides, yet only 37% produce cobamides *de novo*. We were able to predict cobamide structure for 58% of cobamide producers. Additionally, our predictions revealed that 17% of bacteria can salvage cobamide precursors, of which we have defined a new category of bacteria that require early tetrapyrrole precursors to produce cobamides.

## Materials and Methods

### Data set download and filtering

The names, unique identifiers, and metadata for 44,802 publicly available bacterial genomes on the Joint Genome Institute’s Integrated Microbial Genomes with Expert Review database (JGI/IMGer, https://img.jgi.doe.gov/cgi-bin/mer/main.cgi) (Markowitz *et al.*, 2012) classified as “finished” (accessed January 11, 2017) or “permanent draft” (accessed February 23, 2017) were downloaded (Supplementary Table 1, Sheet 1). To assess genome completeness, we searched for 55 single copy gene annotations (Raes *et al.*, 2007; Brown *et al.*, 2015) using the “function profile: genomes vs functions” tool in each genome (Supplementary Table 1, Sheet 4). Completeness was measured first based on the unique number of single copy gene annotation hits (55/55 was best) and second, by the average copy number of the annotations (values closest to 1 were considered most complete) (Supplementary Table 3). We removed 2,776 genomes with fewer than 45 out of 55 unique single copy genes (Supplementary Fig. 1). To filter the remaining genomes to one genome per species, we used name-based matching to create species categories, in which each unique binomial name was considered a single species. The genome with the highest unique single copy gene number and had an average single copy gene number closest to 1 was chosen to represent a species. If both scores were identical the representative genome was chosen at random. For strains with genus assignments, but without species name assignments, we considered each genome to be a species. The list of species was manually curated for species duplicates caused by data entry errors (Supplementary Table 2).

### Detection of cobamide biosynthesis and dependence genes in genomes

Annotations from Enzyme Commission (EC) numbers (http://www.sbcs.qmul.ac.uk/iubmb/enzyme/), Pfam, TIGRFAM, Clusters of Orthologous Groups (COG), and IMG Terms (Cornish-Bowden, 2014; Finn *et al.*, 2016; Haft *et al.*, 2012; Galperin *et al.*, 2015; Markowitz *et al.*, 2012) for cobamide biosynthesis, cobamide-dependent enzymes, and cobamide-independent alternative functions were chosen. These included annotations used by Degnan *et al*. (2014a), but in other cases alternative annotations were chosen to improve specificity of the identified genes (Supplementary Table 4). For example, EC: 4.2.1.30 for glycerol dehydratase identifies both cobamide-dependent and -independent isozymes, so pfam annotations specific to the cobamide-dependent version were used instead. These genes were identified in each genome using the “function profile: genomes vs functions” tool (Jan-May 2017) (Supplementary Table 1, 2 sheet 2). The gene hits were downloaded as a list of gene unique identifiers, gene locus ID, function hit, and genome name (data available upon request).

For genes without functional annotations in the IMGer database, we chose sequences that were genetically or biochemically characterized to use as the query genes in one-way BLASTP (Altschul *et al.*, 1997) against the filtered genomes using the IMGer “gene profile: genomes vs genes” tool, accessed Jan-May 2017 (Supplementary Table 4).

Output files for the cobamide genes were combined into a master file in Microsoft Excel (Supplementary Table 1, 2 sheet 2). This master file was used as input for custom python 2.7 code that interpreted the presence or absence of genes as predicted phenotypes. We used Microsoft Excel and python for further analysis. Genomes were scored for the presence or absence of cobamide-dependent enzymes and alternatives (Supplementary Table 5) based on the annotations in Supplementary Table 4. We then created criteria for seven cobamide biosynthesis phenotypes: very likely cobamide producer, likely cobamide producer, possible cobamide producer, tetrapyrrole precursor salvager, cobinamide (Cbi) salvager, likely non-producer, and very likely non-producer (Supplementary Table 7) and classified genomes accordingly (Supplementary Table 5).

To distinguish putative phenolic lower ligand attachment *arsAB* homologs from other *cobT* homologs that are not known to produce phenolyl cobamides, IMGer entries for all genes that were annotated as *cobT* homologs were downloaded. Tandem *cobT* homologs were defined as those with sequential IMG gene IDs. This list of tandem *cobT* genes was then filtered by size to eliminate genes encoding less than 300 or more than 800 amino acid residues, indicating annotation errors (*cobT* is approximately 350 AA residues) (Supplementary Table 9). The remaining tandem *cobT* homologs were assigned as putative *arsAB* homologs.

To identify the anaerobic benzimidazole biosynthesis genes *bzaABCDEF*, four new hidden Markov model profiles (HMMs) were created and two preexisting ones (TIGR04386 and TIGR04385) were refined. Generally, the process for generating the new HMMs involved performing a Position-Specific Iterated (PSI) BLAST search using previously classified instances of the Bza proteins aligned in Jalview (Altschul *et al.*, 1997; Waterhouse *et al.*, 2009). Due to their similarity, BzaA, BzaB, and BzaF were examined together, as were BzaD and BzaE. To help classify these sequences, Training Set Builder (TSB) was used (Haft and Haft, 2017). All six HMMs have not been assigned TIGRFAM accessions at the time of publication, but will be included in the next TIGRFAM release, and are included as Supplementary Files. Details for each protein are listed in the Supplementary Materials and Methods.

These protein sequences for 10591 of the filtered genomes were queried for each *bza* HMM using hmm3search (HMMER3.1). Hits are only reported above the trusted cutoff defined for each HMM (Supplementary Table 8). A hit for *bzaA* and *bzaB* or *bzaF* indicated that the genome had the potential to produce benzimidazole lower ligands. The specific lower ligand was predicted based on the *bza* genes present (Hazra *et al.*, 2015).

We used BLASTP on IMGer to search for tetrapyrrole precursor biosynthesis genes that appeared to be absent in the 201 species identified as tetrapyrrole precursor salvagers. Query sequences used were the following: *Rhodobacter sphaeroides* HemA (GenPept C49845); *Clostridium saccharobutylicum* DSM 13864 HemA, HemL, HemB, HemC, and HemD (GenBank: AGX44136.1, AGX44131.1, AGX44132.1, AGX44134.1, AGX4133.4, respectively). Since the *C. saccharobutylicum* HemD is a fusion protein with both the UroIII synthase and UroIII methyltransferases domains, we additionally searched for the *Bacillus subtilis* HemD, which only has the UroIII synthase activity (UniProtKB P21248.2). We visually inspected the ORFs near any BLASTP hits in the IMGer genome browser. 180 species remained after this analysis (Supplementary Table 10). Genomes were classified as a particular type of tetrapyrrole precursor salvager only if they were missing all genes upstream of a precursor.

### Strains and growth conditions

*Clostridium scindens* ATCC 35704, *Clostridium sporogenes* ATCC 15579, and *Treponema primitia* ZAS-2 (gift from Jared Leadbetter) were grown anaerobically with and without added 5-aminolevulinic acid (1 mM for *C. sporogenes* and *T. primitia*, 0.5 mM for *C. scindens*).

*Desulfotomaculum reducens* MI-1 (gift from Rizlan Bernier-Latmani), *Listeria monocytogenes* (gift from Daniel Portnoy)*, Blautia hydrogenotrophica* DSM 10507, *Clostridium kluyveri* DSM 555 (gift from Rolf Thauer), and *Clostridium phytofermentans* ISDg (gift from Susan Leschine) were grown anaerobically. Details of the growth conditions are listed in the Supplementary Materials and Methods.

### Corrinoid extraction and analysis

Corrinoid extractions were performed as described (Yi *et al.*, 2012). For corrinoids extracted from 1 L cultures of *C. sporogenes, C. scindens*, and *T. primitia*, high performance liquid chromatrography (HPLC) analysis was performed with an Agilent Series 1200 system (Agilent Technologies, Santa Clara, CA) equipped with a diode array detector with detection wavelengths set at 362 and 525 nm. 50 to 100 µl samples were injected onto an Agilent Eclipse XDB C18 column (5 µm, 4.6 × 150 mm) at 35 °C, with 0.5 ml min^−1^ flow rate. Samples were separated using acidified water and methanol (0.1% formic acid) with a linear gradient of 18% to 30% methanol over 20 min.

For all other bacteria excluding *B. hydrogenotrophica*, extracted corrinoids were analyzed as above, except with a 1.5 ml/min flow rate and a 40°C column. Corrinoids were eluted with the following method: 2% acidified methanol for 2 min, 2%-10% acidified methanol in 0.1 min, and 10-40% acidified methanol over 9 min.

For *B. hydrogenotrophica*, corrinoids were analyzed as above with the following changes. 10 µl samples were injected onto an Agilent Zorbax SB-Aq column (5 µm, 4.6 × 150) with 1 ml/min flow rate at 30°C. The samples were separated with a gradient of 25-34% acidified methanol over 11 minutes, followed by 34-50% over 2 min and 50-75% over 9 minutes.

## Results

### Most bacteria are predicted to have at least one cobamide-dependent enzyme

We surveyed publicly available bacterial genomes for 51 functions involved in cobamide biosynthesis, modification and salvage, as well as 15 cobamide-dependent enzyme families and five cobamide-independent alternative enzymes and pathways. Because most of these genes have been characterized and annotated, we used annotations from existing databases including Enzyme Commission (EC) numbers, Pfam, TIGRFAM, Clusters of Orthologous Groups (COG), and IMG Terms to identify most of these functions and used homology-based methods to identify those for which annotations were unavailable. After filtering 44,802 publicly available bacterial genomes to one genome per species and removing incomplete genomes, we had a working data set of 11,436 species (for details, see Materials and Methods).

Our results indicate that the capability to use cobamides is widespread in bacteria. 86% of species in the filtered data set have at least one of the 15 cobamide-dependent enzyme families shown in Fig. 1 and Supplementary Table 4, and 88% of these species have more than one family (Fig. 3A). This is consistent with previous analyses of smaller data sets (Rodionov *et al.*, 2003; Zhang *et al.*, 2009; Degnan *et al.*, 2014a). The four major phyla in the data set have different distributions of the number of cobamide-dependent enzyme families per genome, with the Proteobacteria and Bacteroidetes having higher mean numbers of enzyme families than the Firmicutes and Actinobacteria (Fig. 3A). The most abundant cobamide-dependent enzymes are involved in core metabolic processes such as methionine synthesis and nucleotide metabolism, whereas processes such as reductive dehalogenation and mercury methylation are less abundant (Fig. 3B, Supplementary Table 5). We also observe phylum-level differences in the relative abundance of cobamide-dependent enzyme families (Fig. 3B), most notably the nearly complete absence of epoxyqueuosine reductase in Actinobacteria. Nonetheless, the cobamide-dependent methionine synthase (MetH) and, to a lesser extent, methylmalonyl-CoA mutase (MCM) and the cobamide-dependent ribonucleotide reductase (RNR), are the most abundant cobamide-dependent enzyme families in all of the four phyla (Fig. 3B).

**Figure 3:**
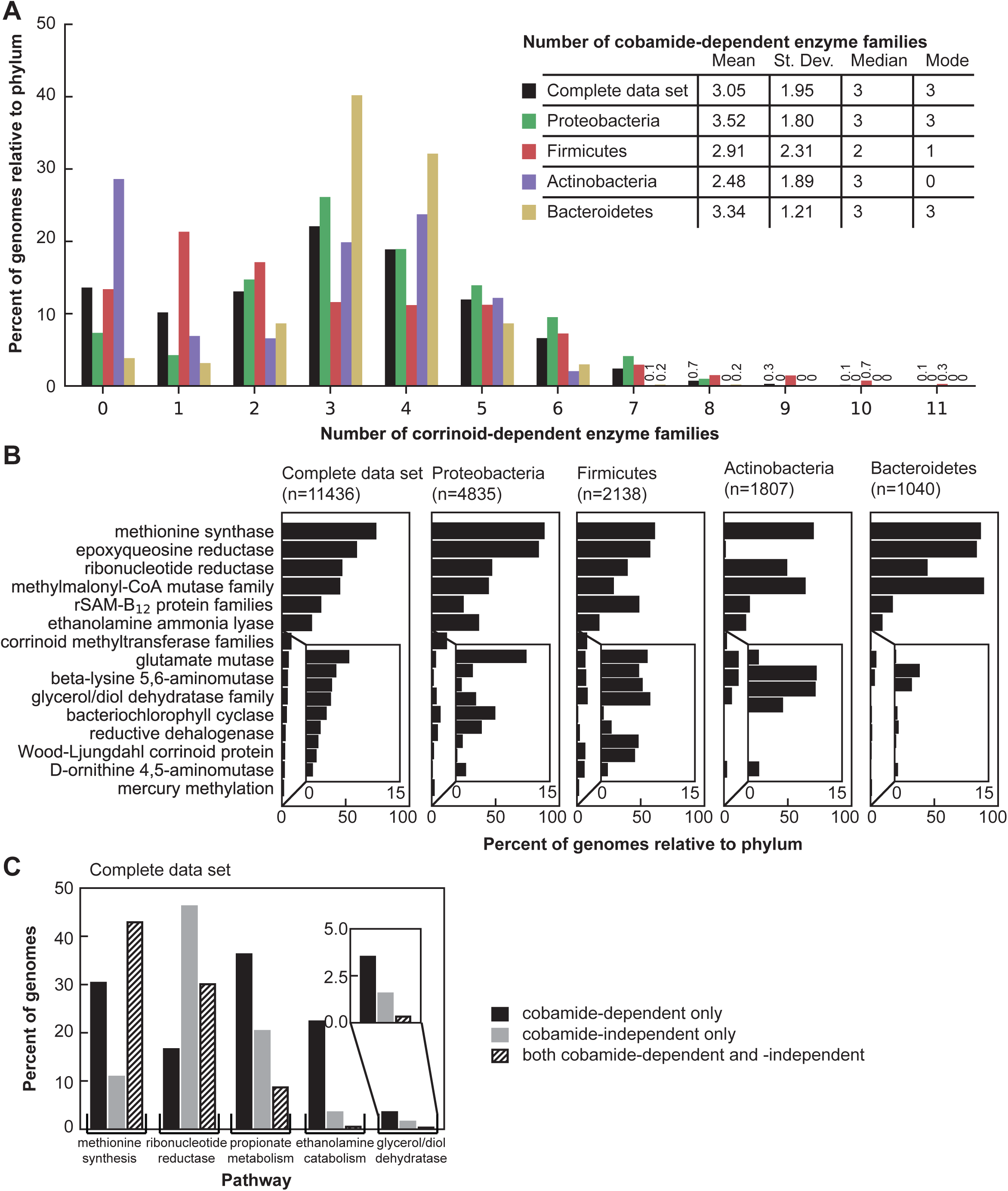
Cobamide dependence in bacteria. **A.** Histogram of the number of cobamide-dependent enzyme families (shown in Fig. 1, Supplementary table 4) per genome in the complete filtered data set and the four most abundant phyla in the data set. The numbers are given for bars with values less than 1%. The inset lists the mean, standard deviation, median, and mode of cobamide-dependent enzyme families for each phylum. **B.** Rank abundance of cobamide-dependent enzyme families in the filtered data set and the four most abundant phyla. The inset shows an expanded view of the nine less abundant functions. **C.** Abundance of five cobamide-dependent processes and cobamide-independent alternatives in the complete filtered data set. Genomes with only the cobamide-dependent, only the cobamide-independent, or both pathways are shown for each process.

For some cobamide-dependent processes, cobamide-independent alternative enzymes or pathways also exist (Fig. 1, right side of arrows). For example, we find that the occurrence of MetH is more common than the cobamide-independent methionine synthase, MetE, but that most bacteria have both enzymes (Fig. 3C). In contrast, cobamide-independent RNRs are found more commonly than the cobamide-dependent versions, and 30% of genomes have both cobamide-dependent and -independent RNRs (Fig. 3C). The cobamide-dependent propionate (which uses MCM), ethanolamine, and glycerol/propanediol metabolisms appear more abundant than the cobamide-independent alternatives (Fig. 3C). However, the abundance of the cobamide-dependent propionate function is overestimated because the MCM category includes mutases for which cobamide-independent versions have not been found. The abundance of the latter two cobamide-independent functions may be underestimated, as they were identified based on similarity to a limited number of sequences. We did not observe dramatic phylum-level differences in the relative abundances of cobamide-dependent and –independent processes (Supplementary Figure 2).

### 37% of bacterial species are predicted to produce cobamides *de novo*

We analyzed the filtered data set to make informed predictions of cobamide biosynthesis to determine the extent of cobamide biosynthesis in bacteria and to identify marker genes predictive of cobamide biosynthesis. A search for genomes containing the complete pathways for anaerobic or aerobic cobamide biosynthesis, as defined in the model bacteria *Salmonella enterica* serovar Typhimurium and *Pseudomonas denitrificans*, respectively (Warren *et al.*, 2002), revealed that few genomes contain all annotations for the complete pathway, but many contain nearly all. Some bacteria that appear to have an incomplete pathway might nonetheless be capable of cobamide biosynthesis because of poor annotation, non-homologous replacement of certain genes (McGoldrick *et al.*, 2005; Gray and Escalante-Semerena, 2010), or functional overlap of some of the enzymes. We therefore relied on experimental data on cobamide biosynthesis in diverse bacteria to inform our predictions, using 63 bacteria that are known to produce cobamides (Table 1, Supplementary Table 6), including 5 tested in this study (Table 1, bold names, Supplementary Figure 3). We identified a core set of eight functions shared by all or all except one of the genomes of cobamide-producing bacteria (Table 1, gray highlight). These core functions include three required for corrin ring biosynthesis: *cbiL*, *cbiF* and *cbiC* in the anaerobic pathway, which are orthologous to *cobI*, *cobM* and *cobH*, respectively, in the aerobic pathway (Table 1, Fig. 2A). An additional five nucleotide loop assembly functions were also highly abundant in these genomes (Table 1).

**Table 1.**
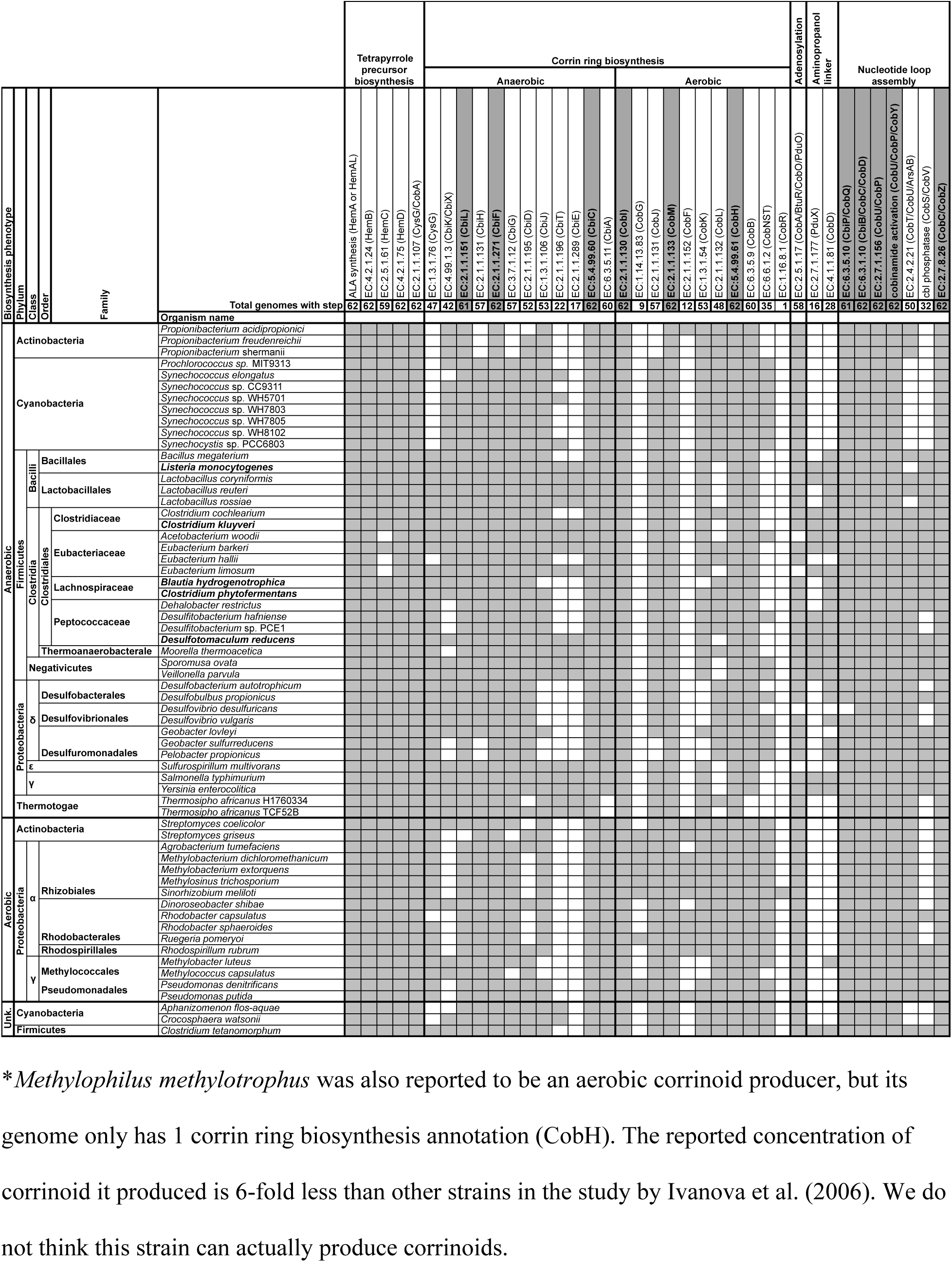
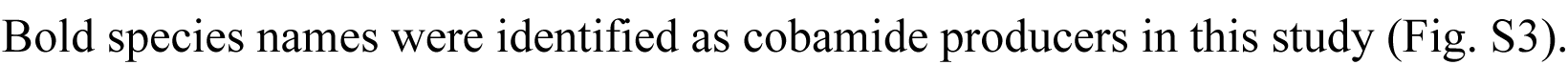
Experimentally-verified cobamide producers and their cobamide biosynthesis annotation content.

Our analysis additionally showed that the anaerobic and aerobic corrin ring biosynthesis pathways cannot be distinguished based on their annotated gene content, presumably because portions of the two pathways share orthologous genes (Table 1; Fig. 2A, dashed lines). Even the cobalt chelatases, *cobNST* and *cbiX/cbiK*, are not exclusive to genomes with the aerobic or anaerobic pathways, respectively (Table 1). Cobalt chelatase annotations are also found in some bacteria that lack most of the corrin ring and nucleotide loop assembly genes, suggesting that there is overlap in annotations with other metal chelatases (Schubert *et al.*, 1999).

We next sought to predict cobamide biosynthesis capability across bacteria by analyzing the filtered genome data set by defining different levels of confidence for predicting cobamide biosynthesis (Supplementary Table 7). Annotations that are absent from the majority of genomes of experimentally verified cobamide producers (*cobR*, *pduX*, and *cobD*) (Table 1, Fig. 2A), as well as one whose role in cobamide biosynthesis has not been determined (*cobW*) (Haas *et al.*, 2009) were excluded from these threshold-based definitions. We did not exclusively use the small set of core functions identified in Table 1 because a correlation between the absence of these genes and lack of cobamide biosynthesis ability has not been established. Using these threshold-based definitions, we predict that 37% of bacteria in the data set have the potential to produce cobamides (Fig. 4, black bars). 49% of species in the data set have at least one cobamide-dependent enzyme but lack a complete cobamide biosynthetic pathway. Genomes in the latter category can be further divided into non-producers, which contain fewer than five corrin ring biosynthesis genes, and precursor salvagers, which contain distinct portions of the pathway (described in a later section). The distribution of cobamide-dependent enzyme families also varies based on predicted cobamide biosynthesis, with predicted cobamide producers having more cobamide-dependent enzyme families per genome than non-producers (Supplementary Figure 4).

**Figure 4:**
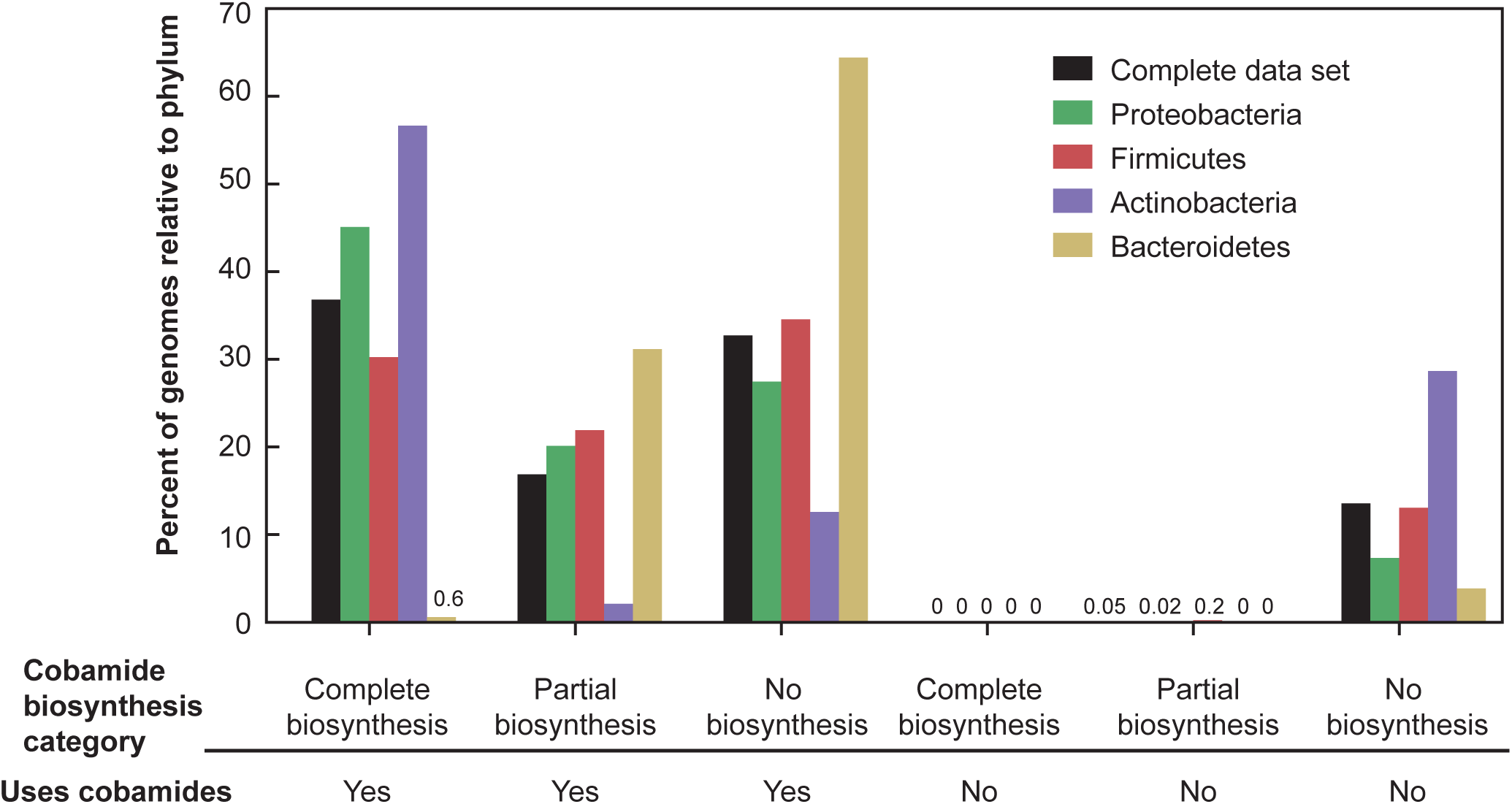
Predicted cobamide biosynthesis phenotypes in the complete filtered data set and the four most abundant phyla in the data set. Genomes were classified into predicted corrinoid biosynthesis phenotypes based on the criteria listed in Supplementary Table 7. The “Partial biosynthesis” category includes cobinamide salvagers and tetrapyrrole precursor auxotrophs. The “Uses cobamides” category is defined as having one or more of the cobamide-dependent enzyme families shown in Figure 1. The numbers are given for bars that are not visible.

To assess whether the core functions could be used as markers, the threshold-based assignments of cobamide biosynthesis were compared to the frequency of the three core corrin ring functions. The presence of each core function alone is largely consistent with the threshold-based assignments, as each is present in 99% of genomes in the producer categories and in less than 1% of the non-producers (Table 2). The presence of two or all three marker functions matches the threshold-based predictions even more closely (Table 2). The corrin ring markers chosen in Table 1 are slightly more predictive of our threshold-based cobamide biosynthesis classifications than *cbiA/cobB* (EC:6.3.5.11/EC:6.3.5.9), a previously selected marker used in environmental DNA analysis (Bertrand *et al.*, 2011); although *cbiA/cobB* was found in 99% of predicted cobamide producers, is it also present in 2.6% of predicted non-producers and 46% of precursor salvagers (Supplementary Table 5).

**Table 2.**
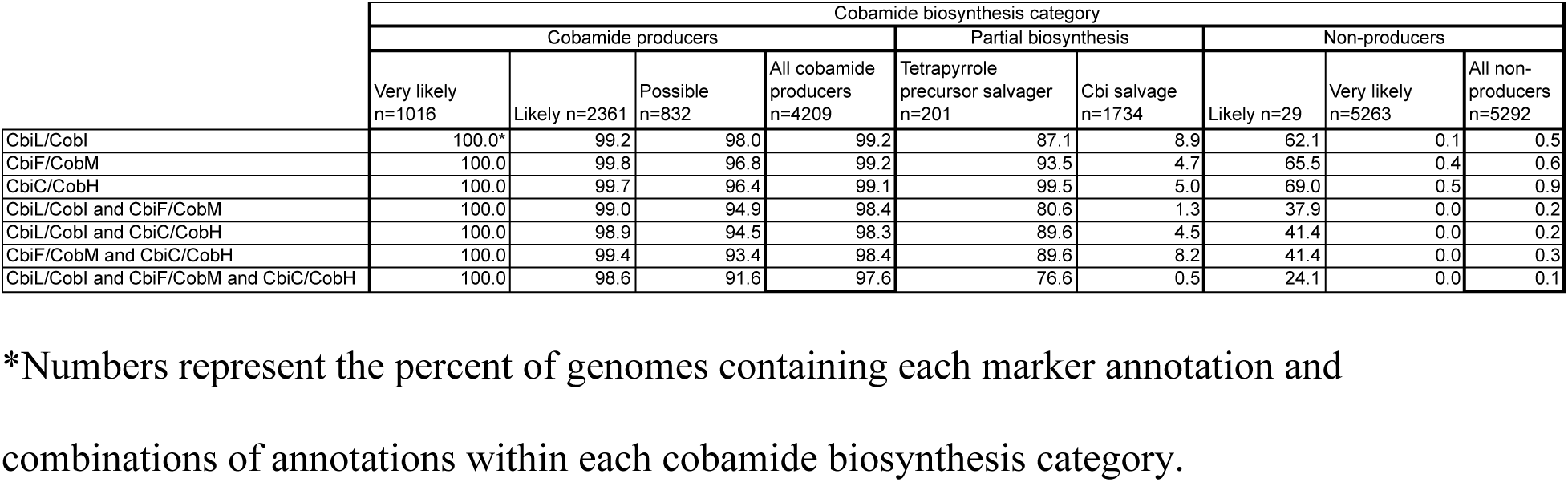
Presence of corrin ring marker annotations in predicted cobamide biosynthesis categories

As with the cobamide-dependent enzyme families, the four major phyla in the data set have major differences in their predicted cobamide biosynthesis phenotypes (Fig. 4). Around half of Actinobacteria (57%) and Proteobacteria (45%) and 30% of Firmicutes are predicted to be cobamide producers. In contrast, only 0.6% of Bacteroidetes are predicted to produce cobamides *de novo*, yet 96% have at least one cobamide-dependent enzyme, suggesting that most members of this phylum must acquire cobamides from other organisms in their environment. In addition, Bacteroidetes have the highest relative proportion of species predicted to salvage Cbi via a partial cobamide biosynthesis pathway, and most of the tetrapyrrole precursor salvagers are Firmicutes (see later section; Supplementary Table 10), whereas very few Actinobacterial species are predicted to salvage precursors (Fig. 4). These divisions reveal potential cobamide and cobamide precursor crossfeeding requirements across phyla.

### Predicting cobamide structure

Lower ligand structure is determined by the intracellular production of lower ligand bases as well as specific features of the lower ligand attachment genes *cobT* or *arsAB* (Campbell *et al.*, 2006; Taga *et al.*, 2007; Gray and Escalante-Semerena, 2007; Hazra *et al.*, 2013; Crofts *et al.*, 2014a; Hazra *et al.*, 2015; Yan *et al.*, 2018; Chan and Escalante-Semerena, 2011). We first defined predictions for the biosynthesis of cobamides containing benzimidazole lower ligands (benzimidazolyl cobamides), based on the presence of genes for the biosynthesis of benzimidazoles. We used the presence of *bluB*, the aerobic synthase for the lower ligand of cobalamin, 5,6-dimethylbenzimidazole (DMB), as a marker for cobalamin production (Campbell *et al.*, 2006; Taga *et al.*, 2007; Hazra *et al.*, 2018) and found it in 25% of genomes in the data set, including those without complete cobamide biosynthesis pathways. *bluB* is most abundant in predicted cobamide-producing bacteria (Fig. 5A), particularly in Proteobacteria (Fig. 5B).

**Figure 5:**
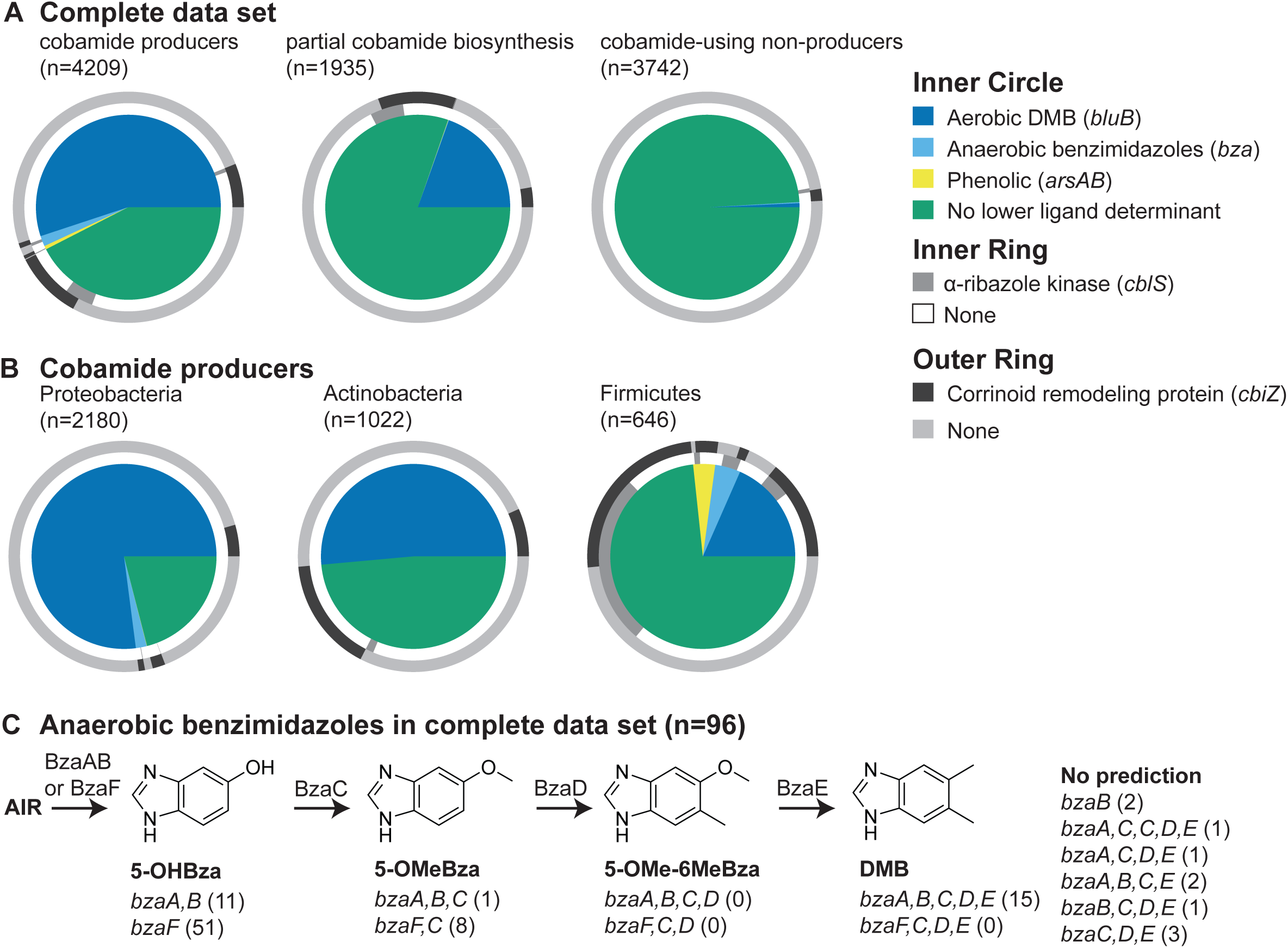
Lower ligand structure predictions. **A, B.** Proportion of genomes containing the indicated lower ligand structure determinants (inner circle), *α*-ribazole salvaging gene (inner ring), and corrinoid remodeling gene (outer ring) in the complete filtered data set separated by cobamide producer category (A) and in cobamide producers separated by phylum (B). **C.** The anaerobic benzimidazole biosynthesis pathway is shown with the functions that catalyze each step above the arrows. The genes required to produce each benzimidazole are shown below each structure, with the number of genomes in the complete filtered data set containing each combination of genes in parentheses. Abbreviations: AIR, aminoimidazole ribotide; 5-OHBza, 5-hydroxybenzimidazole; 5-OMeBza, 5-methoxybenzimidazole; 5-OMe-6-MeBza, 5-methoxy-6-methylbenzimidazole.

Anaerobic biosynthesis of DMB and three other benzimidazoles requires different combinations of the *bza* genes as shown in Figures 2A and 5C (Hazra *et al.*, 2015; Mehta *et al.*, 2015). Because annotations for the majority of the *bza* genes were not available, we developed profile HMMs to search for them (see Supplementary Materials and Methods, Supplementary Files). 96 genomes contain one or more *bza* genes, and 88 of these contain either *bzaF* or both *bzaA* and *bzaB*, the first step necessary for the anaerobic biosynthesis of all four benzimidazoles (Fig. 5C, Supplementary Table 8). As seen with *bluB*, anaerobic benzimidazole biosynthesis genes are highly enriched in cobamide producers (Fig. 5A). Examining the set of *bza* genes in each genome allowed us to predict the structures of cobamides produced in 86 out of the 96 genomes (Fig. 5C). Based on the frequency of *bluB* and the *bza* genes, 24% of bacteria are predicted to produce cobalamin, the cobamide required by humans.

To predict the biosynthesis of phenolyl cobamides, we searched for genomes containing two adjacent *cobT* annotations, since the *cobT* homologs *arsA* and *arsB*, which together are necessary for activation of phenolic compounds for incorporation into a cobamide, are encoded in tandem (Chan and Escalante-Semerena, 2011). Using this definition, *arsAB* was found in only 27 species, and is almost entirely restricted to the class Negativicutes in the phylum Firmicutes, which are the only bacteria reported to produce phenolyl cobamides (Stupperich and Eisinger, 1989; Men *et al.*, 2014b) (Fig. 5A, B, Supplementary Table 9).

42% of predicted cobamide producers in the data set do not have any of the benzimidazole biosynthesis or phenolic attachment genes (Fig. 5A). However, bacteria that have the α-ribazole kinase CblS (Fig. 5A, B, inner rings) and the transporter CblT (not included) are predicted to use activated forms of lower ligand bases found in the environment (Fig. 2A, α-ribazole salvaging); we found CblS in 363 species (3.2%), mostly in the Firmicutes phylum (Fig. 5 A, B, inner rings) (Gray and Escalante-Semerena, 2010; Mattes and Escalante-Semerena, 2017). A higher proportion of bacteria, 1,041 species (9.1%), have a CbiZ annotation (Fig. 5A, B, outer rings), an amidohydrolase that cleaves the nucleotide loop, allowing cells to rebuild a cobamide with a different lower ligand (Woodson and Escalante-Semerena, 2004) (Fig. 2A, corrinoid remodeling). CbiZ is found in genomes of predicted cobamide producers and Cbi auxotrophs (see following section) (Fig. 5A), as expected based on experimental studies (Gray and Escalante-Semerena, 2009a, 2009b; Men *et al.*, 2014a; Yi *et al.*, 2012). The reliance of some bacteria on exogenous lower ligands or α-ribazoles produced by other organisms precludes prediction of cobamide structure in all cases.

### 17% of bacteria have partial cobamide biosynthetic pathways

Our analysis of the cobamide biosynthesis pathway revealed two categories of genomes that lack some or most genes in the pathway, but retain contiguous portions of the pathway. Genomes in one category, the Cbi (cobinamide) auxotrophs (15% of genomes), contain the nucleotide loop assembly steps but lack all or most of the corrin ring biosynthesis functions. As demonstrated in *Escherichia coli* (Di Girolamo and Bradbeer, 1976), *Thermotoga lettingae* (Butzin *et al.*, 2013), and *Dehalococcoides mccartyi* (Yi *et al.*, 2012), and predicted in human gut microbes (Degnan *et al.*, 2014a), Cbi auxotrophs can take up the late intermediate Cbi, assemble the nucleotide loop and attach a lower ligand in a process called Cbi salvaging.

We observed an additional 201 genomes (1.7%) that lack one or more initial steps in tetrapyrrole precursor biosynthesis but have complete corrin ring biosynthesis and nucleotide loop assembly pathways, primarily in the Firmicutes (Supplementary Table 7). After searching these genomes manually for genes missing from the pathway, we designated 180 of these species as tetrapyrrole precursor salvagers, a new classification of cobamide intermediate auxotrophs (Fig. 6A, Supplementary Table 10). These organisms are predicted to produce cobamides only when provided with a tetrapyrrole precursor or a later intermediate in the pathway.

**Figure 6:**
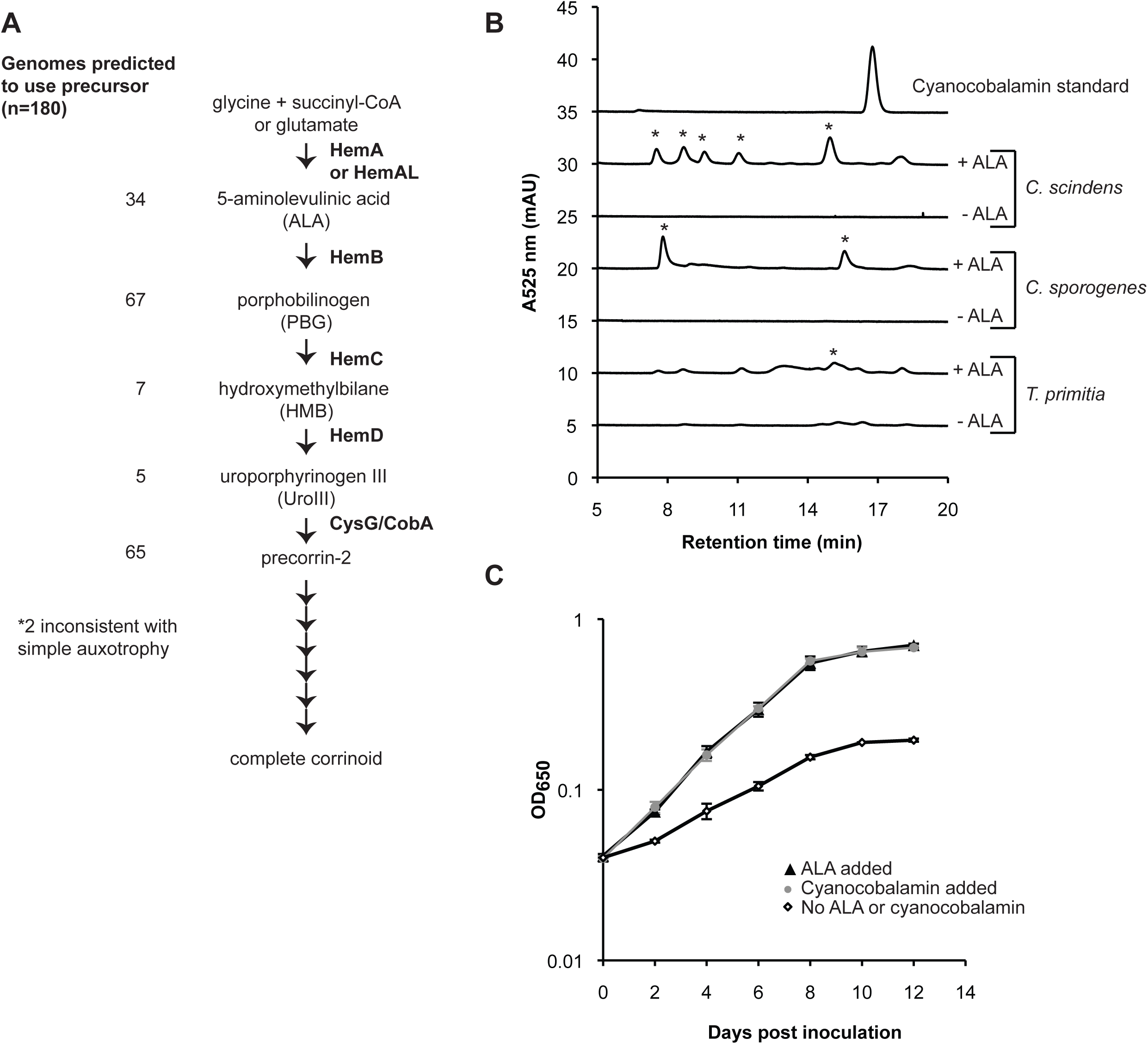
Characterization of putative tetrapyrrole precursor salvagers. **A.** Early steps in cobamide biosynthesis. The functions that catalyze each step are indicated to the right of each arrow. The number of genomes in the complete filtered data set in each tetrapyrrole precursor salvage category is on the left. Two genomes had cobamide biosynthesis pathways inconsistent with simple auxotrophy. Specific genomes are listed in Supplementary Table 10. **B.** HPLC analysis of corrinoid extracts from *Clostridium scindens*, *Clostridium sporogenes*, and *Treponema primitia* grown with and without added ALA. A cyanocobalamin standard (10 μM) is shown for comparison. Asterisks denote peaks with UV-Vis spectra consistent with that of a corrinoid. **C.** *T. primitia* ZAS-2 growth in 4YACo medium with and without added cyanocobalamin or ALA.

### Experimental validation of ALA dependence

The identification of putative tetrapyrrole precursor salvagers suggests either that these bacteria are capable of taking up a tetrapyrrole precursor from the environment to produce a cobamide or that they synthesize the precursors through a novel pathway. We therefore tested three putative tetrapyrrole precursor salvagers for their ability to produce corrinoids (cobamides and other corrin ring-containing compounds) in the presence and absence of a tetrapyrrole precursor. *Clostridium scindens* and *Clostridium sporogenes*, which are predicted to require 5-aminolevulinic acid (ALA), produced corrinoids in defined media only when ALA was supplied, suggesting that they do not have a novel ALA biosynthesis pathway (Fig. 6B). We tested an additional predicted ALA salvager, the termite gut bacterium *Treponema primitia* ZAS-2, for which a defined medium has not been developed. When cultured in medium containing yeast autolysate, *T. primitia* produced trace amounts of corrinoids, and corrinoid production was increased by supplementing this medium with ALA (Fig. 6B). The ability of *T. primitia* to use externally supplied ALA was further shown by its increased growth rate and cell density at stationary phase when either cobalamin or ALA was added (Fig. 6C). Together, these results support the hypothesis that predicted ALA salvagers synthesize cobamides by taking up ALA from the environment.

## Discussion

Vitamin B_12_ and other cobamides have long been appreciated as a required nutrient for humans, bacteria, and other organisms due to their critical function as enzyme cofactors. Prior to this work, the extent of cobamide biosynthesis and dependence across different bacterial taxa had not been investigated. The availability of tens of thousands of genome sequences afforded us the opportunity to conduct a comprehensive analysis of cobamide metabolism across over 11,000 bacterial genomes. This analysis gives an overview of cobamide dependence and cobamide biosynthesis across bacteria, allowing for the generation of hypotheses for cobamide and cobamide precursor crossfeeding in bacterial communities. Our work shows that cobamide use is much more widespread than cobamide biosynthesis, consistent with the majority of previous studies of smaller data sets (Rodionov *et al.*, 2003; Zhang *et al.*, 2009; Degnan *et al.*, 2014a). The prevalence of cobamide-dependent enzymes in bacteria, coupled with the relative paucity of *de novo* cobamide producers, underscores the ubiquity of both cobamide-dependent metabolism and cobamide crossfeeding in microbial communities. Here, we additionally find that cobamide production and use are unevenly distributed across the major phyla represented in the data set, identify bacteria dependent on cobamide precursors, and predict cobamide structure. These results underscore the widespread nutritional dependence of bacteria.

The most abundant types of cobamide-dependent enzymes in our data set are methionine synthase, epoxyqueuosine reductase, RNR, and MCM. For all of these enzymes, cobamide-independent alternative enzymes or pathways exist. (Note that the newly discovered alternative to epoxyqueuosine reductase, QueH (Zallot *et al.*, 2017), was not included in our analysis.) The prevalence of cobamide-dependent enzymes for which cobamide-independent counterparts exist, particularly in the same genome, suggests that cobamide-dependent enzymes confer distinct advantages. This view is supported by the observations that MetE is sensitive to stress and has a 100-fold lower turnover number than MetH (Hondorp and Matthews, 2004; Xie *et al.*, 2013; Gonzalez *et al.*, 1992) and that cobamide-independent RNRs are limited in the oxygen concentrations in which they are active (Taga and Walker, 2010; Fontecave, 1998)

In our analysis of cobamide biosynthesis, it was not possible to use a single definition of the complete *de novo* cobamide biosynthesis pathway across all bacterial genomes because of divergence in sequence and function. The use of experimental data gives confidence to our predictions and allowed identification of marker genes for cobamide biosynthesis. Nevertheless, our predictions likely overestimate the extent of cobamide biosynthesis *in situ*, as genome predictions do not account for differences in gene expression. For example, cobamide production in *S. typhimurium* is repressed in environments containing oxygen or lacking propanediol (Roth *et al.*, 1996), and cobamide biosynthesis operons are commonly subjected to negative regulation by riboswitches (Nahvi, 2004; Rodionov *et al.*, 2003). The abundance of cobamide importers (Rodionov *et al.*, 2003; Zhang *et al.*, 2009; Degnan *et al.*, 2014a), even in bacteria capable of cobamide biosynthesis, reinforces the possibility that many bacteria may repress expression of cobamide biosynthesis genes in favor of cobamide uptake in some environments.

A comparison of genomes containing one or more cobamide-dependent functions to those with none revealed an absence of bacteria that produce cobamides but do not use them. This finding suggests that altruistic bacteria that produce cobamides exclusively for others do not exist. Metabolically coupled organisms that crossfeed cobalamin in exchange for another nutrient have been described in the mutualistic relationships between algae and cobalamin-producing bacteria (Croft *et al.*, 2005; Kazamia *et al.*, 2012), yet it remains unclear if such intimate partnerships are widespread. Notably, our results show that cobamide biosynthesis is unevenly distributed across bacteria, with Actinobacteria enriched in and Bacteroidetes lacking in *de novo* cobamide biosynthesis. Such phylogenetic comparisons can be used to make crude predictions of cobamide crossfeeding relationships among different taxa.

The reliance of many bacteria on cobamide crossfeeding, coupled with the fact that structurally different cobamides are not functionally equivalent in bacteria (Yan *et al.*, 2012; Mok and Taga, 2013; Degnan *et al.*, 2014a; Helliwell *et al.*, 2016; Keller *et al.*, 2018), underscores the importance of cobamide structure in microbial interactions. The structural differences in cobamides are almost exclusively limited to variations in the lower ligand (Fig. 2B). Additional variation in the nucleotide loop was not considered here because of the absence of genes specific to norcobamide biosynthesis (Kräutler *et al.*, 2003; Keller *et al.*, 2016). We were able to predict lower ligand structure for 58% of predicted cobamide producers. The remaining bacteria may produce purinyl cobamides, which are abundant in some bacterial taxa and microbial communities (Helliwell *et al.*, 2016; Allen and Stabler, 2008). Further analysis of substrate specificity in CobT and other lower ligand attachment enzymes could lead to improved strategies for predicting purinyl cobamide production, as some CobT homologs appear to segregate into different clades based on lower ligand structure (Hazra *et al.*, 2013; Crofts *et al.*, 2013; Yan *et al.*, 2018). The presence of free benzimidazoles and *α*-ribazoles in microbial communities (Crofts *et al.*, 2014b; Johnson *et al.*, 2016; Wienhausen *et al.*, 2017) and the ability of bacteria to take up and incorporate these compounds into cobamides (Anderson *et al.*, 2008; Mok and Taga, 2013; Keller *et al.*, 2013; Crofts *et al.*, 2013) suggest that it will not be possible to predict the structures of cobamides produced by all bacteria *in situ* solely from genomic analysis.

We predict that 32% of cobamide-dependent bacteria are unable to synthesize cobamides, attach a preferred lower ligand to Cbi, or remodel corrinoids. This group of bacteria must take up cobamides from their environment for use in their cobamide-dependent metabolisms. Given the variable use of structurally different cobamides by different bacteria, the availability of specific cobamides is likely critical to bacteria that are unable to synthesize cobamides or alter their structure. The availability of preferred cobamides may limit the range of environments that these organisms can inhabit. Variation in the abundance of different cobamides has been observed in different environments. For example, in a TCE-contaminated groundwater enrichment culture, 5-hydroxybenzimidazolyl cobamide and *p*-cresolyl cobamide were the most abundant cobamides (Men *et al.*, 2014b), compared to cobalamin in bovine rumen (Girard *et al.*, 2009) and 2-methyladeninyl cobamide in human stool (Allen and Stabler, 2008). One strategy for acquiring preferred cobamides could be selective cobamide import, as suggested by the ability of two cobamide transporters in *Bacteroides thetaiotaomicron* to distinguish between different cobamides (Degnan *et al.*, 2014a).

Dependence on biosynthetic precursors has been observed or predicted for amino acids, nucleotides, and the cofactors thiamin and folate (Sloan and Moran, 2012; Kilstrup *et al.*, 2005; Paerl *et al.*, 2016; de Crécy-Lagard *et al.*, 2007). Here, we describe genomic evidence for dependence on cobamide precursors, namely Cbi or tetrapyrrole precursors. The prevalence of Cbi-salvaging bacteria suggests that it is common for bacteria to fulfill their cobamide requirements by importing Cbi from the environment and assembling the nucleotide loop intracellularly. Consistent with this, Cbi represented up to 9% of total corrinoids in TCE-contaminated groundwater enrichments (Men *et al.*, 2014b), and represented up to 12.8% of the total corrinoids detected in human stool samples (Allen and Stabler, 2008).

Our analysis defined five types of tetrapyrrole precursor salvagers and experimentally verified the ALA salvager phenotype in three species. Bacteria that lack tetrapyrrole precursor biosynthesis genes but contain the remainder of the cobamide biosynthetic pathway were overlooked in previous genomic studies of cobamide biosynthesis that considered only the corrin ring biosynthesis and nucleotide loop assembly the portions of the pathway (Rodionov *et al.*, 2003; Zhang *et al.*, 2009; Magnúsdóttir *et al.*, 2015; Helliwell *et al.*, 2016). Tetrapyrrole precursors have been detected in biological samples, suggesting that they are available for uptake in some environments. For example, UroIII was detected in human stool (Dobriner, 1937; Watson *et al.*, 1945) and ALA has been found in swine manure extract (Kanto *et al.*, 2013). Although we confirmed experimentally the ALA dependence phenotype, we were unable to detect ALA in several biological samples using a standard chemical assay or bioassay, suggesting either that ALA is not freely available in these environments or is present at concentrations lower than the 100 nM detection limit of these assays (data not shown). Based on the ecosystem assignment information available for 48% of the genomes, 78% of tetrapyrrole precursor salvagers are categorized as host-associated bacteria compared to 41% in the complete filtered dataset. One interpretation of this finding is that tetrapyrrole precursors are provided by the host, either from host cells that produce them as intermediates in heme biosynthesis (Sangwan and O’Brian, 1991; Lyell *et al.*, 2017) or, for gut-associated microbes, as part of the host’s diet. Alternatively, these precursors may be provided by other microbes, as was observed in a coculture of *Fibrobacter* species (Qi *et al.*, 2008). Genome analysis suggests that Candidatus *Hodgkina cicadicola*, a predicted Uroporphyrinogen III (UroIII) salvager (McCutcheon *et al.*, 2009), may acquire a tetrapyrrole precursor from its insect host or other endosymbionts to be able to provide methionine for itself and its host via the cobamide-dependent methionine synthase. 17% of cobamide-requiring human gut bacteria lacked genes to make UroIII de novo from glutamate, suggesting they could be UroIII salvagers (Degnan *et al.*, 2014a).

Nutritional dependence is nearly universal in bacteria. Auxotrophy for B vitamins, amino acids, and nucleic acids is so common that these nutrients are standard components of bacterial growth media. We speculate that the availability of cobamides in the environment, coupled with the relative metabolic cost of cobamide biosynthesis, has driven selection for loss of the cobamide biosynthesis pathway (Morris *et al.*, 2012). The large number of genomes with partial cobamide biosynthesis pathways, namely in the “possible cobamide biosynthesis”, “likely non-producer”, and “Cbi salvager” classifications, suggests that some of these genomes are in the process of losing the cobamide biosynthesis pathway. At the same time, evidence for horizontal acquisition of the cobamide biosynthesis pathway suggests an adaptive advantage for nutritional independence for some bacteria (Morita *et al.*, 2008; Lawrence and Roth, 1996). Such advantages could include early colonization of an environmental niche, ability to synthesize cobamides with lower ligands that are not commonly available, or association with hosts that do not produce cobamides. The analysis of the genomic potential of bacteria for cobamide use and production presented here could provide a foundation for future studies of the evolution and ecology of cobamide interdependence.

## Acknowledgments

This work was supported by NIH grant AI117984 and a Hellman Family Faculty Fund award to M.E.T., a Grace Kase Graduate Fellowship to A.N.S., and an NSF Graduate Fellowship to E.C.S, and NSF grant No. 1458808 to the J. Craig Venter Institute.

We thank Alison Smith, Johan Kudahl, Jared Leadbetter, and members of the Taga lab for helpful advice. We thank Olga Sokolovskaya, Sebastian Gude, Zachary Hallberg, and Alexa Nicolas for critical reading for this manuscript.

We thank Jared Leadbetter, Rizlan Bernier-Latmani, Daniel Portnoy, Rolf Thauer, and Susan Leschine for providing bacterial strains.

## Contributions

A.N.S., E.C.S., and A.W.H. performed the bioinformatic analysis. S.N.J. and D.R.H. created the BzaABCDEF HMMs. K.C.M., E.C.S., and A.N.S. performed growth assays and corrinoid extractions.

